# NF-κB-Driven HIV-1 Gene Expression in Human Cells Is Independent of Poly(ADP-ribose) polymerase-1 Function

**DOI:** 10.1101/2025.03.10.642491

**Authors:** Denisse A. Gutierrez, Manuel Llano

**Author notes:** Corresponding author. Address correspondence to: Manuel Llano. Department of Biological Sciences, The University of Texas at El Paso, 500 W. University Ave. El Paso, TX 79968, USA.

## Abstract

The cellular enzyme poly (ADP-ribose) polymerase-1 (PARP-1) is required for NF-κB to activate inflammatory and immune response gene expression. NF-κB is also an important transcription factor in HIV-1 gene expression during active replication and latency reactivation. Therefore, enhancing NF-κB signaling is an alternative for HIV-1 latency reactivation, but significant systemic side effects related to the NF-κB role in inflammatory and immune responses are predictable. To verify this prediction, we determined whether PARP-1 is required in NF-κB-dependent HIV-1 gene expression in a human CD4+ T lymphoblastoid cell line (SUP-T1) and HEK 293T cells. Our findings indicated that PARP-1 knockout does not impair HIV-1 infection or gene expression. Specifically, NF-κB-dependent HIV-1 gene expression was not impaired by PARP-1 deficiency, highlighting an important transcriptional regulatory difference between HIV-1 and inflammatory and immune activation genes. Our findings define a negligible role of PARP-1 in HIV-1 gene expression, suggesting that PARP-1 antagonism could ameliorate the expected inflammatory response with latency-reactivating agents that act through the NF-κB signaling pathway.

**Importance:** PARP-1 is required for NF-κB to activate the expression of inflammatory and immune response genes. NF-κB is also an important transcription factor in HIV-1 gene expression during active replication and latency reactivation. Enhancing NF-κB signaling is expected to cause HIV-1 latency reactivation, but significant systemic side effects related to the NF-κB role in inflammatory and immune responses are predictable. The role of PARP-1 in NF-κB-mediated activation of HIV-1 gene expression and in viral infection has not been determined in the context of HIV-1 infection of CD4+ T cells. Our data indicate that PARP-1 is dispensable for NF-κB-mediated activation of HIV-1 gene expression in a human CD4+ T lymphoblastoid cell line. These findings suggest that the pharmacological antagonism of PARP-1 could diminish the inflammatory effects of latency-reactivating agents that activate NF-κB signaling without impairing their effect on HIV-1 gene expression.

## INTRODUCTION

HIV cure requires the removal of the latently infected reservoir (1, 2). Transcriptional regulation of proviruses is central to any strategy of eliminating the functional reservoir (3). Among the different signaling pathways regulating HIV-1 transcription, the NF-κB pathway is very relevant in latency establishment and reversal (4, 5). However, because of the essential role of NF-κB regulation in the expression of inflammatory and other immune response genes (6), interventions that trigger HIV-1 transcription in an NF-κB-dependent manner are expected to cause severe inflammation and immune activation (4, 7–14)

Poly (ADP-ribose) polymerase-1 (PARP-1), an enzyme implicated in multiple cellular processes, including the regulation of transcription (15, 16), modulates NF-κB activity through both enzymatic and non-enzymatic mechanisms. This functional interaction regulates the transcription of a wide variety of host genes implicated in inflammation and immune activation (17–22). PARP-1 is the most enzymatically active member of the PARP family, promoting the transfer of ADP ribose molecules from NAD+ to acceptor proteins or to an existing poly (ADP-ribose) (PAR) chain (23). This post-translational modification alters the function of target proteins by changing their subcellular localization, molecular interactions, and enzymatic activities. Additionally, catalytic-independent functions have been demonstrated to mediate the PARP-1 biological activities (16, 23–26).

PARP-1 stimulates NF-κB signaling through its coactivator function in response to inflammatory stimuli, and via the atypical NF-κB signaling pathway activated by DNA damage (27). As a coactivator, PARP-1 promotes the interaction of NF-κB with the basal transcription machinery (17, 18, 28), whereas, in response to DNA damage, PARP-1 facilitates the nuclear localization of NF-κB (4, 6, 28). Additionally, through PARylation, PARP-1 enhances the activity of NF-κB (12, 29).

In contrast to the well-established role of PARP-1 in the transcriptional regulation of inflammation and immune activation genes, its function in HIV-1 gene expression is a matter of debate. Different functions of PARP-1 suggest its implication in HIV-1 gene expression (18, 30, 31). At the level of transcriptional initiation, PARP-1 could modulate the activity of several transcription factors or the chromatin structure at the viral promoter (3). PARP-1 has been reported to regulate the activity of NF-κB, AP-1, Sp1, and NFAT (19, 32), which are implicated in the transcription of HIV-1 (3). In particular, NF-κB and Sp1 are crucial for viral transcription, while AP-1 and NFAT play more modulatory roles. NF-κB is sufficient to activate LTR transcription and its activity is potentiated by Sp1 and AP-1 (3). Furthermore, PARP-1 has been reported to enhance Tat activity in an LTR reporter in HeLa cells (33).

In correlation with a positive role of PARP-1 in the activity of NF-κB in the context of the HIV-1 promoter, PARP-1 inhibition has been reported to decrease HIV mRNA levels and virus production in the chronically infected human pro-monocyte U1 cell line stimulated with the NF-κB activator Phorbol 12-myristate 13-acetate (PMA) (30). Additionally, transient PARP-1-KD in Jurkat and HeLa cells, and PARP-1 inhibition in human monocyte-derived macrophages reduced HIV-1 LTR reporter activation triggered by compounds that activate NF-κB signaling (18, 30, 31, 34). In contrast to this positive role in NF-κB-mediated HIV-1 LTR transcription, Vpr, which facilitates HIV-1 replication in T cells and is required for optimal infection of human monocyte-derived macrophages, has been reported to retain PARP-1 in the cytosol, impairing NF-κB-mediated transcription of a non-viral promoter (35).

To better understand the role of PARP-1 in NF-κB-dependent HIV-1 gene expression, we investigated the impact of NF-κB signaling on HIV-1 proviral gene expression in PARP-1-knockout (KO) human CD4+ T cell line (SUP-T1) and HEK 293T cells. Our results demonstrate that PARP-1 deficiency does not impair HIV-1 infection or gene expression and is not essential for NF-κB-mediated HIV-1 transcription. Interestingly, PARP-1 KO significantly disrupted NF-κB-driven activation of the promoter of the inflammatory gene inducible nitric oxide synthase (iNOS). These findings suggest that targeting PARP-1 may reduce the inflammatory response associated with the use of latency-reactivating agents that activate the NF-κB signaling pathway.

## RESULTS

### HIV-1 infection is not impaired in PARP-1-KO human CD4+ T lymphoblastoid cells

Several reports indicate that PARP-1 deficiency does not affect the infectivity of VSV-G pseudotyped, single-round infection HIV-1 (36–38), suggesting that PARP-1 does not affect HIV-1 gene expression. Importantly, in only one of these publications (37), experiments were conducted in cells of a histological origin relevant to HIV-1 infection *in vivo*, meaning in a human CD4+ T cell line (SUP-T1). However, an important caveat in this research was that the cells studied were only partially deficient in PARP-1. To use a more robust cellular model, we evaluated the role of PARP-1 in HIV-1 gene expression, and in particular in NF-κB regulation of the HIV-1 LTR, in PARP-1 knockout cells derived from the human CD4+ T cell line SUP-T1, and their backcomplemented counterpart (39).

PARP-1 KO and BC SUP-T1 cells (Fig. 1a) were infected with a single-round infection, VSV-G pseudotyped HIV-1_NL4-3_ and luciferase levels were measured four days later. This reporter virus is mutated in the *env*, *nef* and *vpr* genes, and expresses LTR-driven luciferase from the *nef* slot (Hluc) (40, 41). In correspondence with previous findings (37, 39), PARP-1 levels did not significantly influence HIV-1 transgene expression (Fig. 1b). KO cells expressed luciferase at 1.1-fold (clone 1) and 0.8-fold (clone 2) of the levels observed in the corresponding BC clones, indicating that PARP-1 has no significant effect on HIV-1 infection or gene expression.

**Figure 1.**
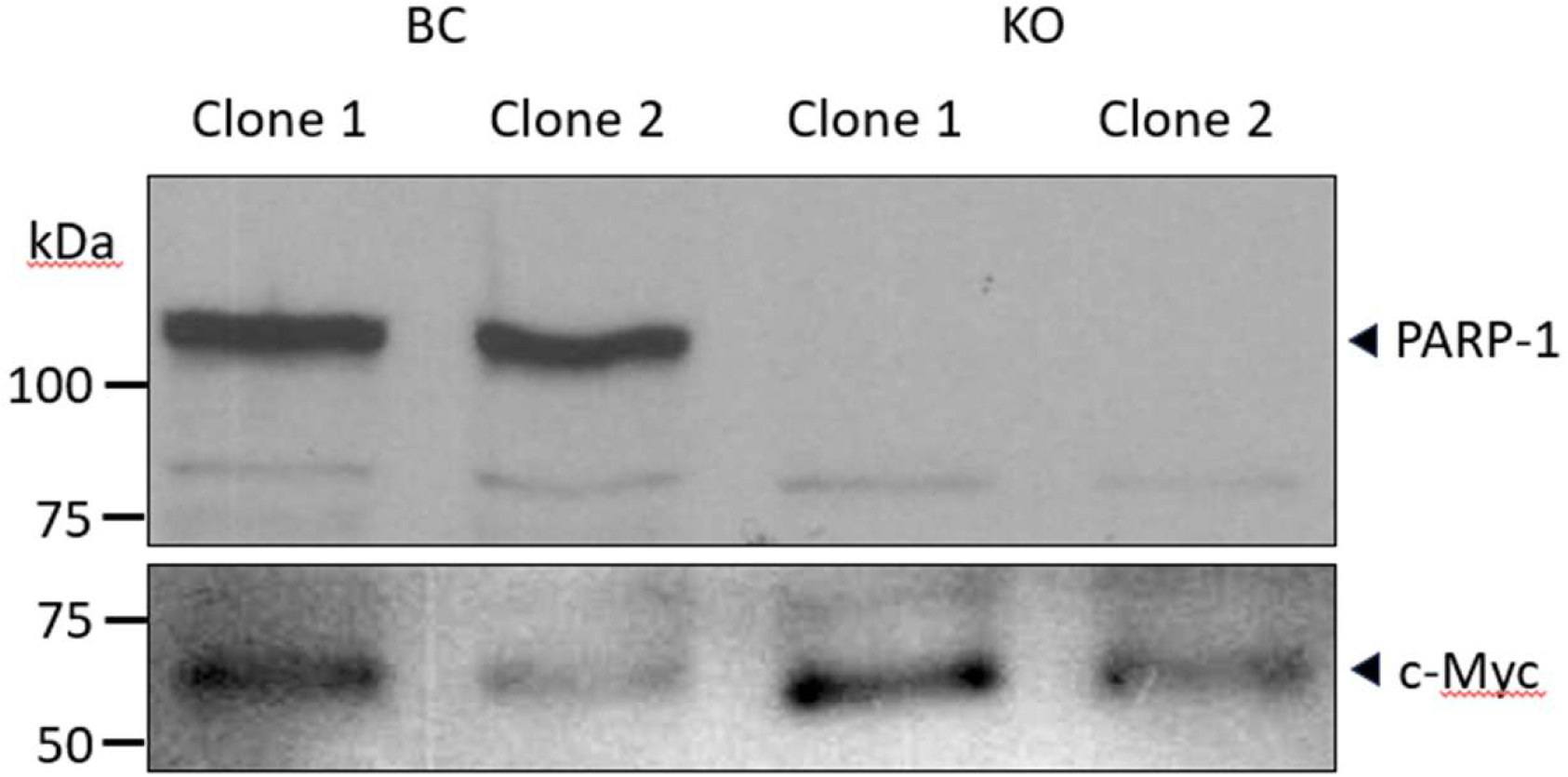
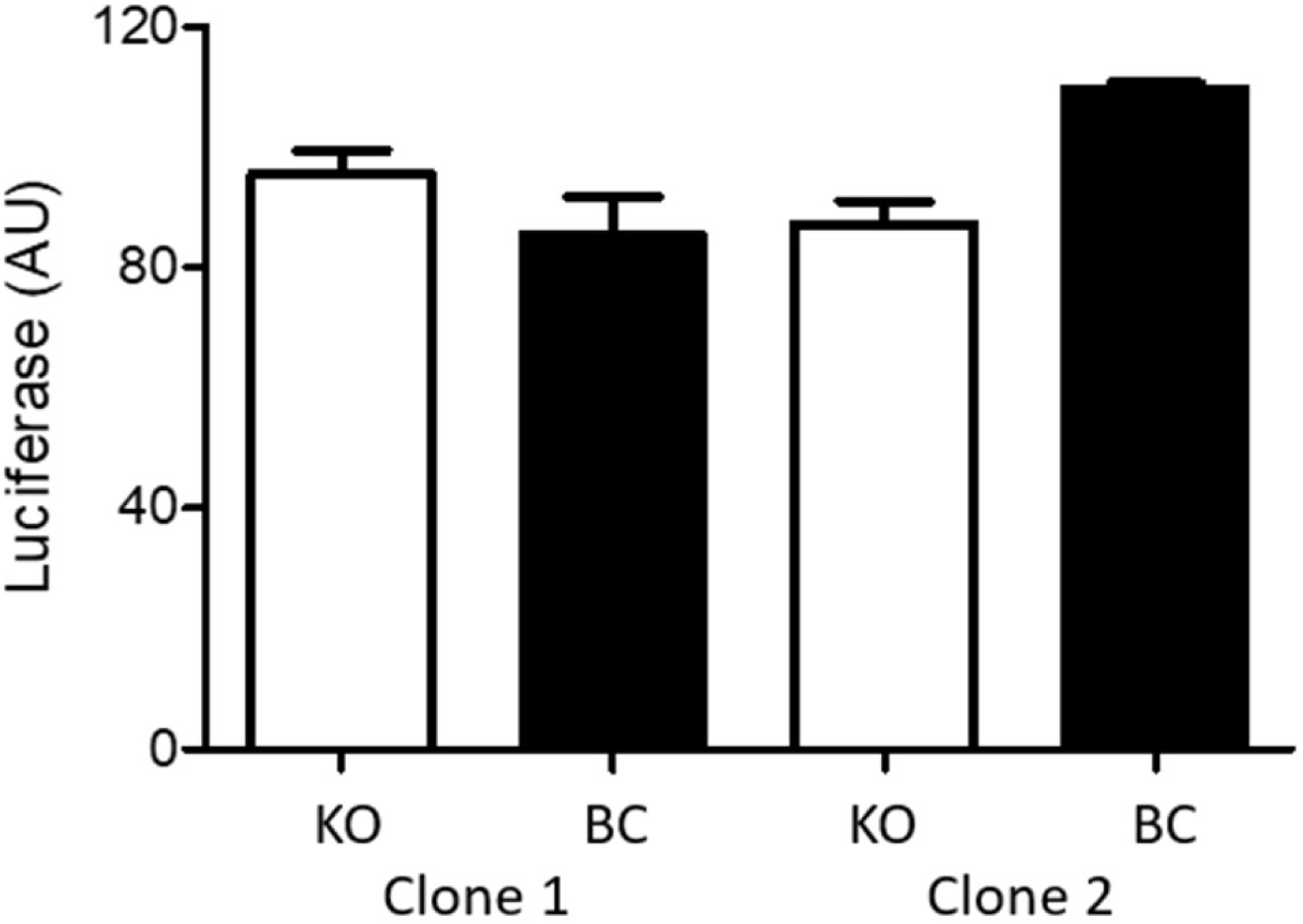
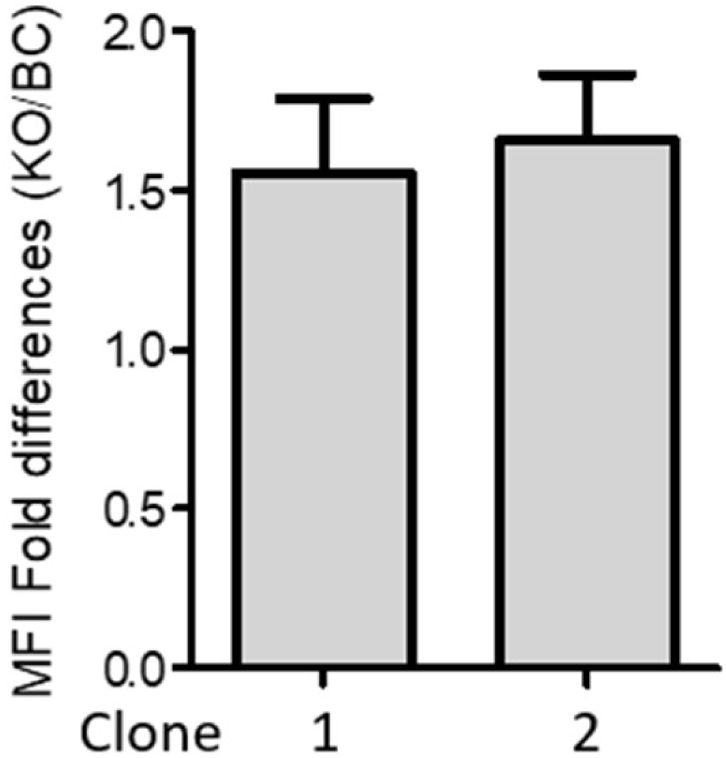
Effect of PARP-1 deficiency on HIV-1 infection. (**a**) Immunoblot analysis of PARP-1 levels in SUP-T1-derived cell lines. Detection of c-Myc was used as a loading control. (**b**) Luciferase levels in cells infected with Hluc. (**c**) eGFP levels in cells infected with NLENG1-ES-IRES. Data in (**b**) and (**c**) correspond to one experiment done in triplicate that is representative of more than three independent experiments performed in triplicate.

We also evaluated the effect of PARP-1 deficiency on the infection and gene expression of an HIV-1_NL4-3_-derivative that expresses eGFP from the viral promoter. PARP-1-KO and -BC clones were infected with this virus and analyzed by FACS four days later (Fig. 1c). eGFP mean fluorescence intensity was 1.5 +/- 0.5 and 1.6 +/- 0.6 higher in PARP-KO clones 1 and 2, respectively, than in their corresponding BC cell lines, demonstrating further that PARP-1 is dispensable for LTR-driven transcription.

### PARP-1 is dispensable in NF-κB-dependent HIV-1 gene expression

PARP-1 is required in NF-κB-dependent expression of multiple pro-inflammatory genes (19, 27, 42, 43) and genes activated by CD3/CD28 signaling in lymphocytes (19, 32). In contrast, PARP-1 seems to be dispensable for the transcriptional activity of the HIV-1 LTR (Fig. 1), which is an NF-κB-responsive promoter. This potential dichotomy is important since the undesired inflammatory response is a relevant side effect of HIV-1 latency reactivation strategies targeting NF-κB.

To assess the potential dispensability of PARP-1 in NF-κB-dependent HIV-1 LTR-driven gene expression, Hluc-infected PARP-1 knockout (KO) and BC SUP-T1 cells were cultured for three weeks to eliminate unintegrated HIV-1 cDNA. The cells were then stimulated with the NF-κB activators tumor necrosis factor-alpha (TNF-α, 10 ng/mL) and phorbol 12-myristate 13-acetate (PMA, 40 and 10 ng/mL). Three days post-stimulation, luciferase expression was measured to evaluate transcriptional activity.

TNF-α (Fig. 2a) equivalently increased luciferase expression in both PARP-1 KO and BC SUP-T1 infected cells. In clone 1, TNF-α increased luciferase 5.7 (KO cells) and 7.4 folds (BC cells). Similarly, in clone 2 the increase was 8.5- and 10-fold in KO and BC cells, respectively. Therefore, activation was only 1.3 (clone 1) and 1.2 (clone 1) higher in BC than in KO SUPT1 cells.

**Figure 2.**
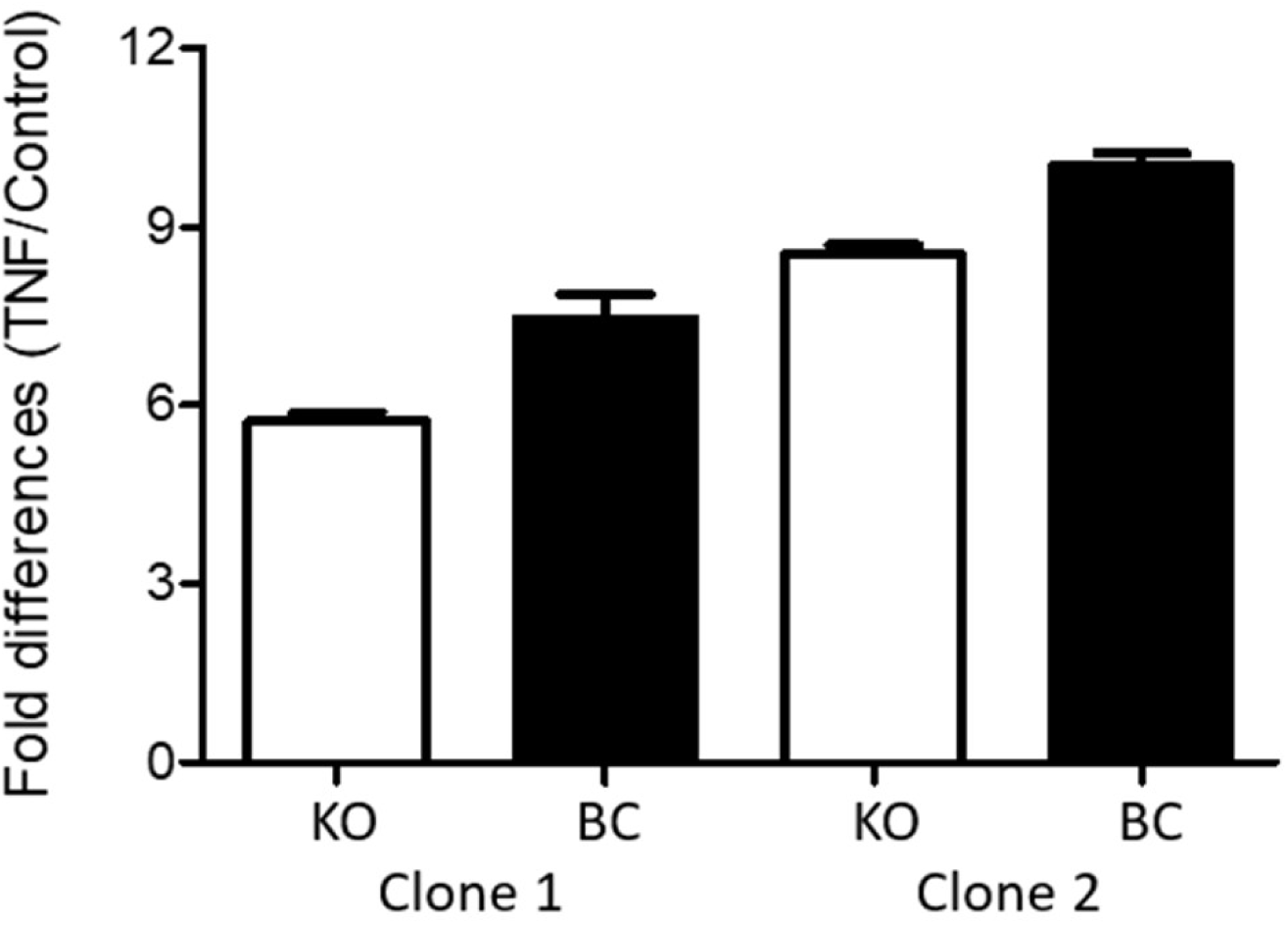
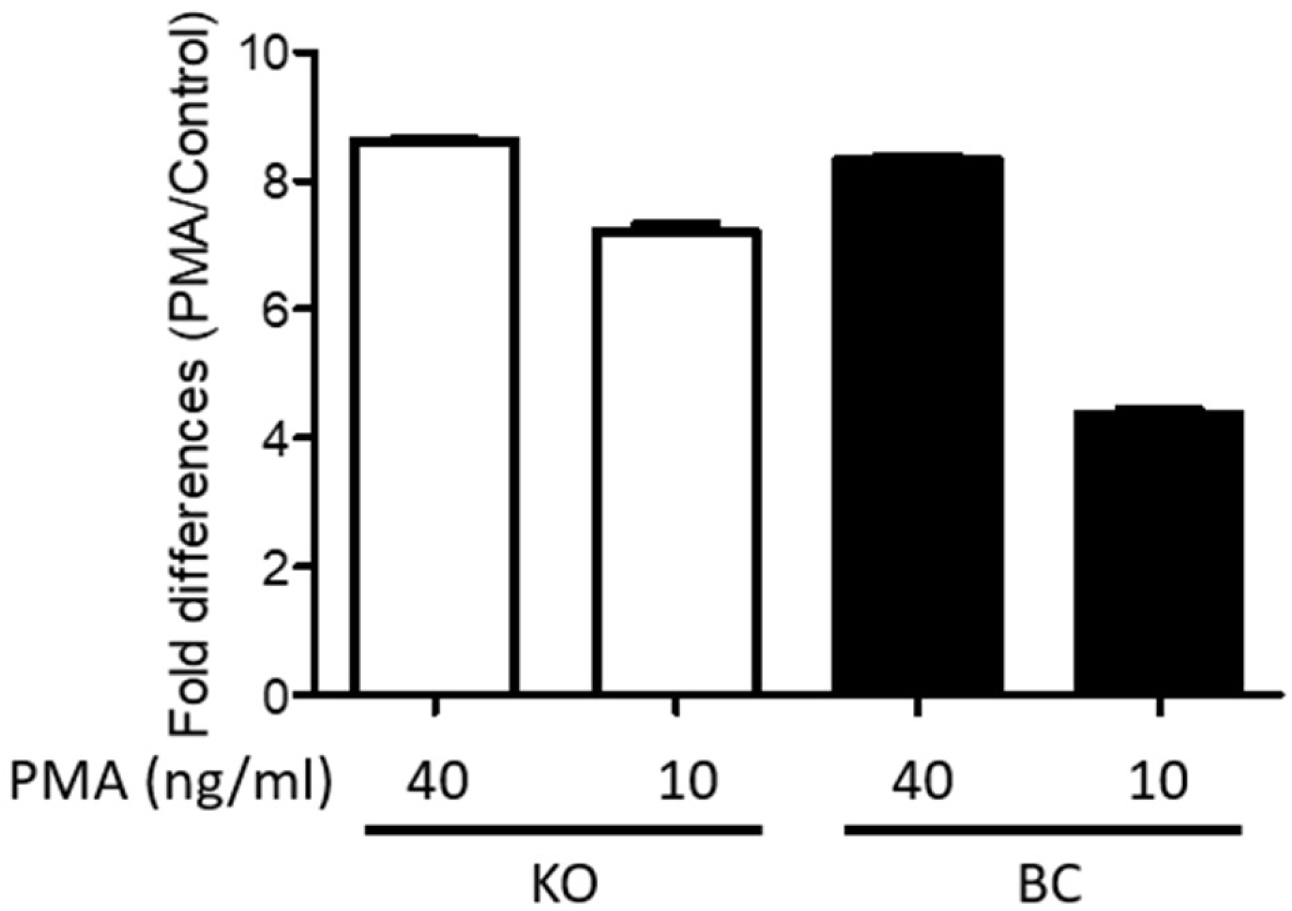
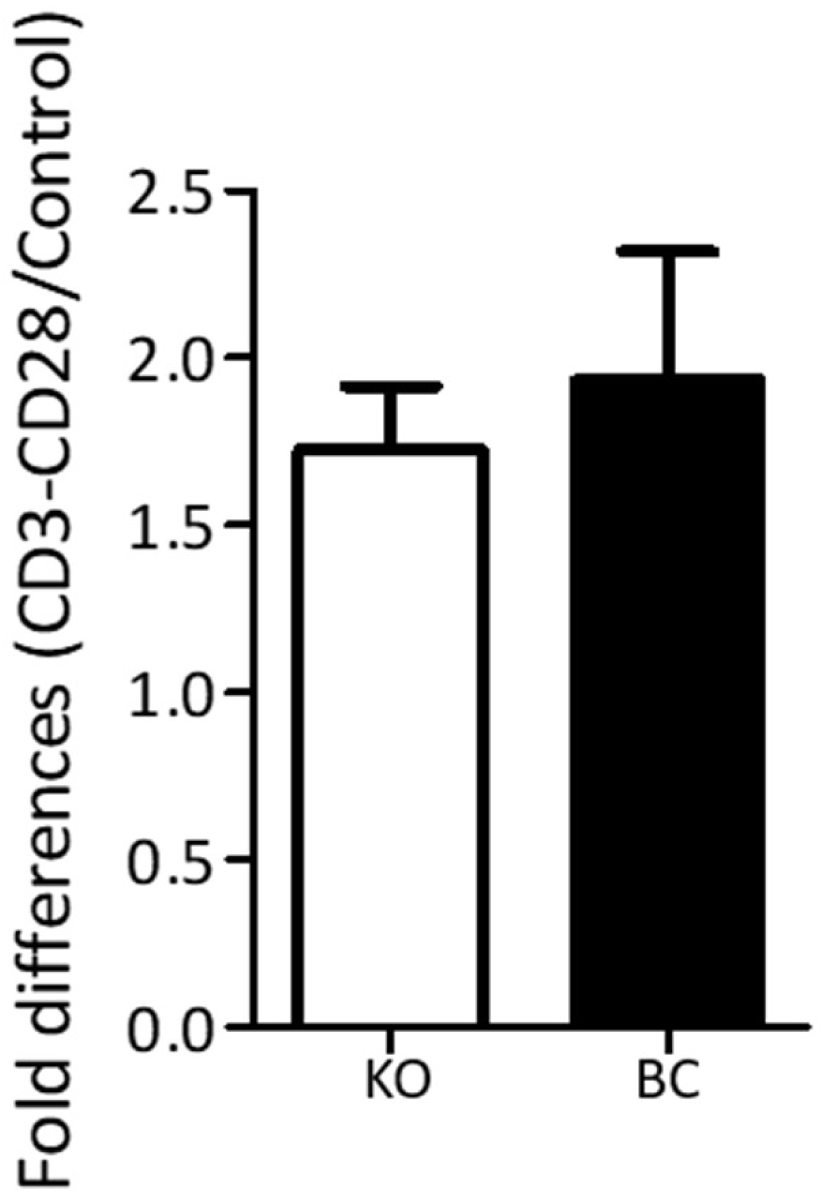
Impact of PARP-1 on HIV-1 gene expression mediated by different transcriptional activators. Hluc-infected PARP-1 KO and BC cells were treated with vehicle (control) or several transcriptional activators, and luciferase levels were determined. Control luciferase levels were used for normalization. (**a**) TNF-α (10 ng/ml), (**b**) PMA (40 and 10 ng/ml), and (**c**) Anti-CD3/CD28 beads. Data in Figure 2 correspond to one experiment done in triplicate that is representative of two independent experiments performed in triplicate.

PMA similarly activated the HIV-1 LTR in PARP-1 KO and BC SUPT1 cells (Fig. 2b). At 40 ng/ml, PMA caused 8.6- and 8.3-fold activation in clone 1 KO and BC cells, respectively. Meanwhile, at 10 ng/ml PMA activated the HIV-1 LTR by 7.2-fold and 4.4-fold in KO and BC SUP-T1 cells, respectively. These results reaffirming that PARP-1 is not required for PMA activation of the HIV-1 LTR promoter.

HIV-infected PARP-1 KO and -BC SUP-T1 cells were also stimulated by anti-CD3/-CD28 crosslinking that activates NFκB, AP-1, and NFAT signaling pathways (19, 32). CD3/CD28 stimulation activated HIV-1 gene expression by 1.7- and 1.9-fold in PARP-1 KO and BC SUP-T1 cells (Fig. 2c). These findings also indicated a negligible role of PARP-1 in the regulation of the HIV-1 promoter through these transcription factors.

TNF-α is expected to stimulate the HIV-1 LTR promoter via NF-κB signaling (44). To verify the implication of this mechanism, PARP-1 KO and BC SUPT1 cells were infected with Hluc, and 4 days later the cells were stimulated for 3 days with TNF-α (10 ng/ml) in the presence or no of BAY-11-7082 (BAY). This compound inhibits the TNF-α-induced phosphorylation of IκB-α, preventing NF-κB nuclear translocation (45).

To exclude treatment toxicity, we first evaluated the effect of 3-day BAY treatment on the viability of SUP-T1 cells, measured as ATP levels. At 10 µM and 5 µM the compound was very toxic, and cell viability dropped to 2% +/- 4.4% and 57% +/- 12.6%, respectively, the viability of control cells treated with DMSO. However, at 3 µM and 2 µM, cell viability was 80% +/- 10.4% and 83% +/-13.5%, respectively, the viability of DMSO-treated cells. Therefore, we used BAY at 3 µM in subsequent studies.

As shown before, basal levels of luciferase were similar in non-treated PARP-1 KO and BC SUPT1 clones 1 (Fig.3a I) and 2 (Fig.3a II), and these values were used for normalization of the luciferase levels found in the corresponding cells upon TNF-α stimulation. Notably, luciferase expression was upregulated by TNF-α stimulation in both PARP-1 KO (6.4 and 1.9 folds, clones 1 and 2) and BC (4.3 and 1.6 folds, clones 1 and 2) SUPT1 cells (Fig.3a). Furthermore, BAY blocked the stimulatory effect of TNF-α with equivalent potency in both PARP-1 KO (2.1 and 2 folds, clones 1 and 2) and BC (2.5 and 3.7 folds, clones 1 and 2) SUPT1 cells (Fig. 3a I - II). These findings indicated that NF-κB mediates TNF-α-induced activation of the HIV-1 promoter, regardless of PARP-1 cellular levels.

**Figure 3.**
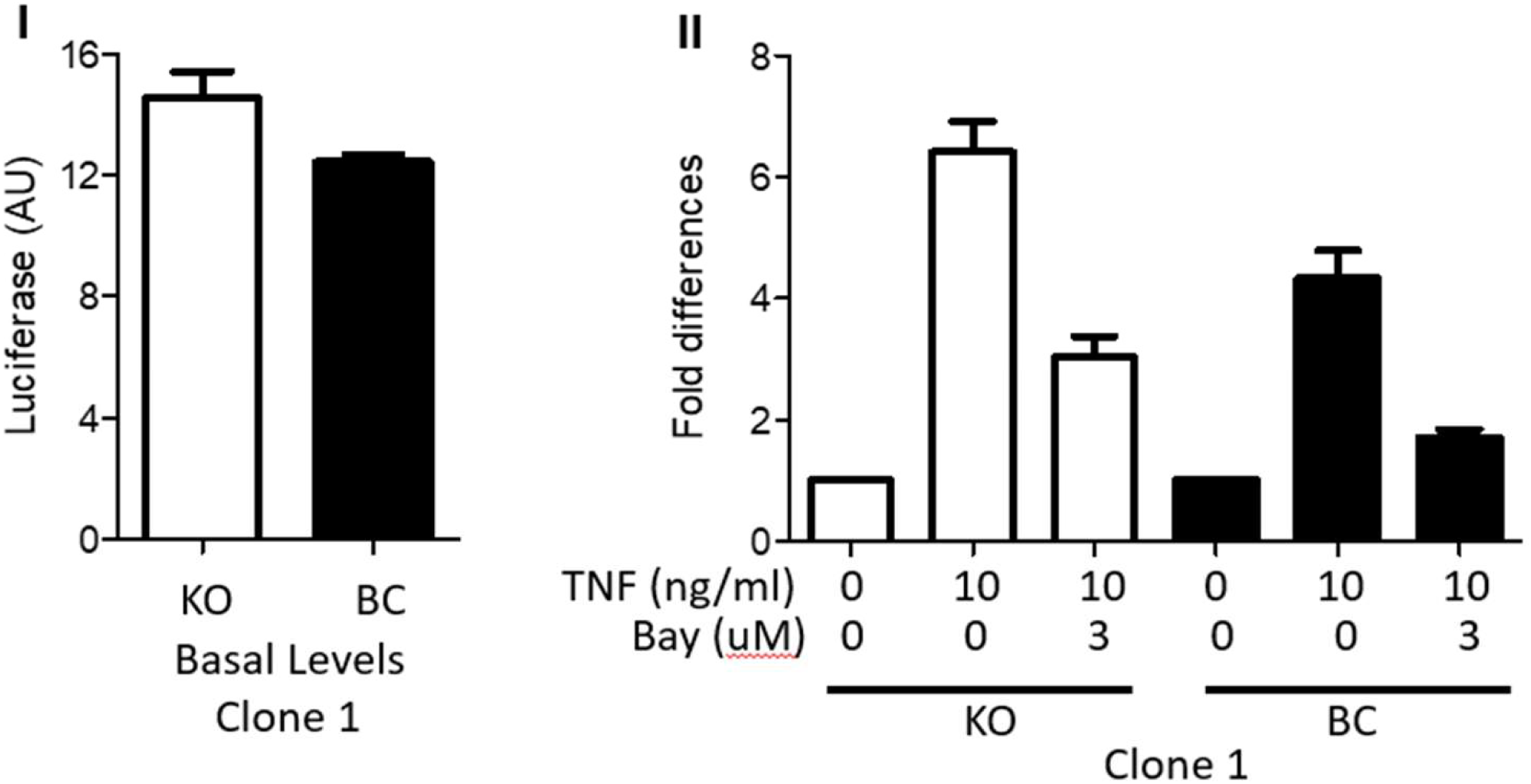
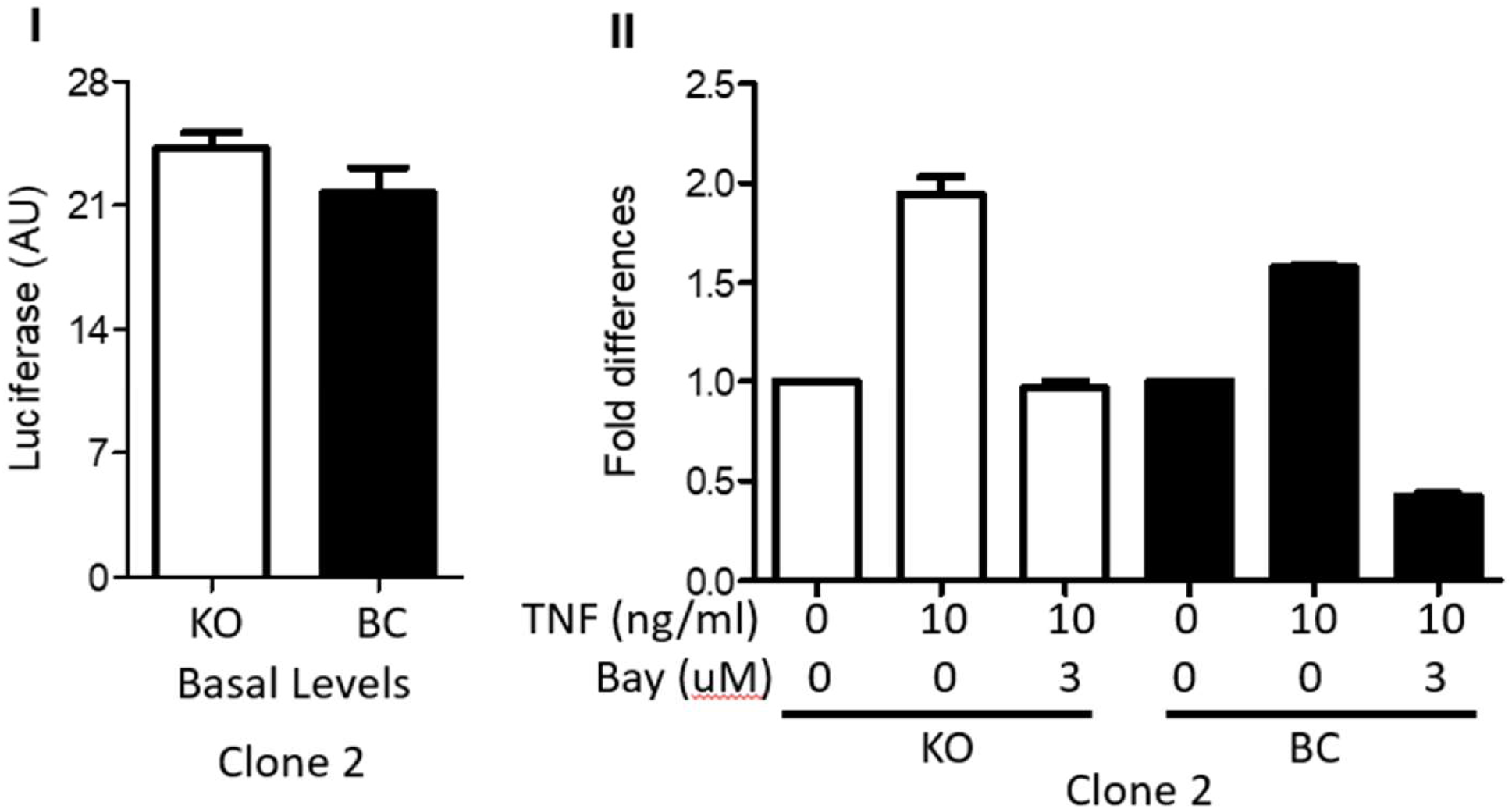
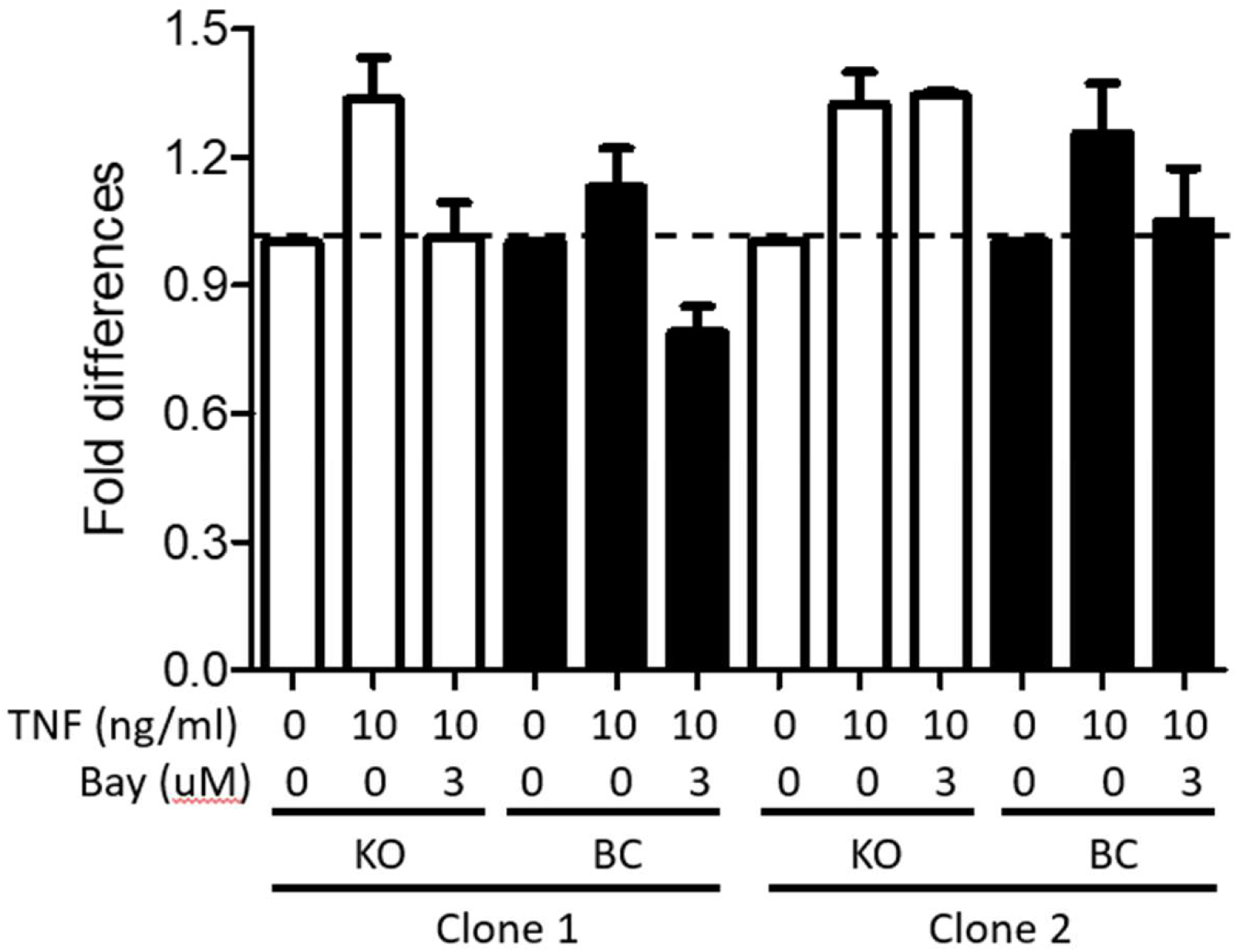
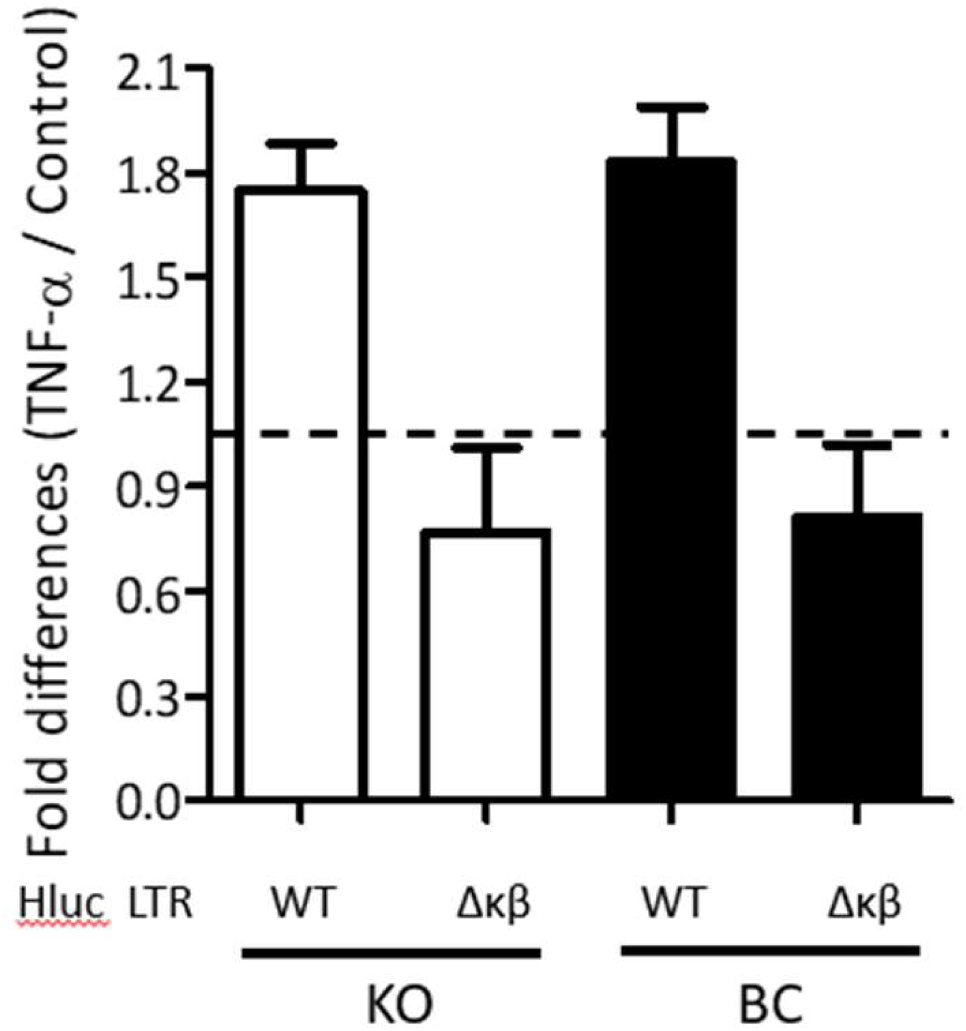
Effect of TNF-α stimulation on HIV-1 gene expression in cells expressing or no PARP-1. (**a** and **b**) (**I**) Basal levels of luciferase in PARP-1 KO and BC SUP-T1 cells infected with Hluc (clone 1, panel **a** and clone 2, panel **b**). (**II**) Cells characterized in panel **I** were treated with TNF-α alone or in the presence of BAY-11-7082 (BAY), and luciferase levels were determined. Basal luciferase levels (**I**) were used for normalization in (**II**). (**c**) ATP levels (cell viability) in the cells analyzed in panels (**a** and **b**), dotted line indicates no change. (**d**) TNF-α-induced luciferase expression in PARP-1 KO and BC cells infected with Hluc or Hluc_Δκβ_. Luciferase levels in vehicle-treated cells (control) were used for data normalization. Data in Figure 3 correspond to one experiment done in triplicate that is representative of three independent experiments performed in triplicate.

The viability of the cells studied in figure 3a was determined by measuring their ATP levels. The basal ATP levels of untreated cells were used to normalize the ATP measurements in their corresponding treated counterparts. Data in Fig. 3b indicate no important differences in cell viability. ATP levels only decreased, to 79% of control values, in PARP-1 BC clone 1 following treatment with TNF-α and BAY. These results confirmed that the reduction in HIV-1 transgene expression induced by BAY was not attributable to cell toxicity.

To further demonstrate that in these cells TNF-α induced HIV-1 gene expression through NF-κB signaling, PARP-1 KO and BC SUP-T1 cells were infected with Hluc carrying an NF-κB mutant (Hluc_Δκβ_) or wild-type LTR. Hluc_Δκβ_ lacks the two copies of the enhancer κB elements implicated in NF-κB binding (46). Cells were stimulated or not with TNF-α at day four post-infection, and three days later; luciferase levels were determined. As shown before (Fig. 2a and 3a), TNF-α activated luciferase expression with a similar magnitude in PARP-1 KO (1.7 folds) and BC (1.8 folds) SUPT1 cells (Fig. 3c). However, TNF-α failed to activate LTR-driven transcription (< 1-fold) in cells infected with Hluc_Δκβ_, independently of the PARP-1 levels in these cells (Fig. 3c). Therefore, TNF-α stimulated HIV-1 gene expression in PARP-1 KO cells in an NF-κB-dependent manner.

### PARP-1 is required in the NF-κB-dependent expression of the inflammatory gene inducible nitric-oxide synthase (iNOS)

As mentioned above, PARP-1 is required for NF-κB-dependent transcriptional activation of inflammatory genes, such as iNOS (47, 48). Therefore, as a control, we evaluated the requirement of PARP-1 in NF-κB-dependent transcriptional activation of the iNOS promoter. To this end, we used PARP-1 KO and BC HEK 293T cells (39) due to their high transfection efficiency. PARP-1 KO and BC HEK 293T cells (Fig. 4a) were co-transfected with a CMV-driven β-galactosidase expression plasmid (transfection control) along with a plasmid encoding a luciferase reporter whose expression is driven by the iNOS promoter wild-type (iNOSp WT) or a mutant promoter lacking the NF-κB-binding site (iNOSpΔNFκB) (49). Two days after transfection, the cells were divided into different groups and treated with vehicle (basal conditions), TNF-α, and TNF-α + BAY. Twenty-four hours later, luciferase and β-galactosidase activity were measured. Luciferase values were normalized to β-galactosidase activity to account for transfection efficiency.

**Figure 4.**
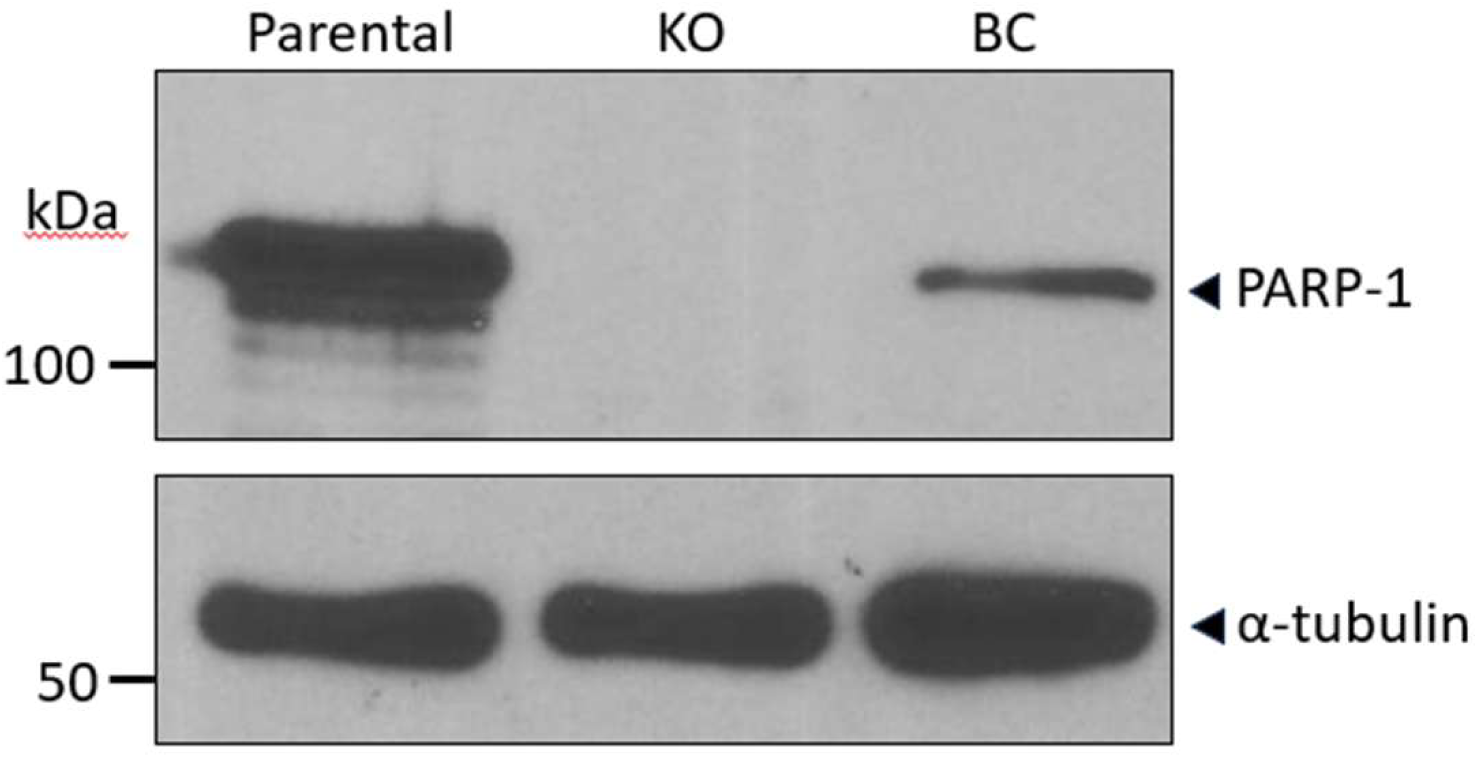
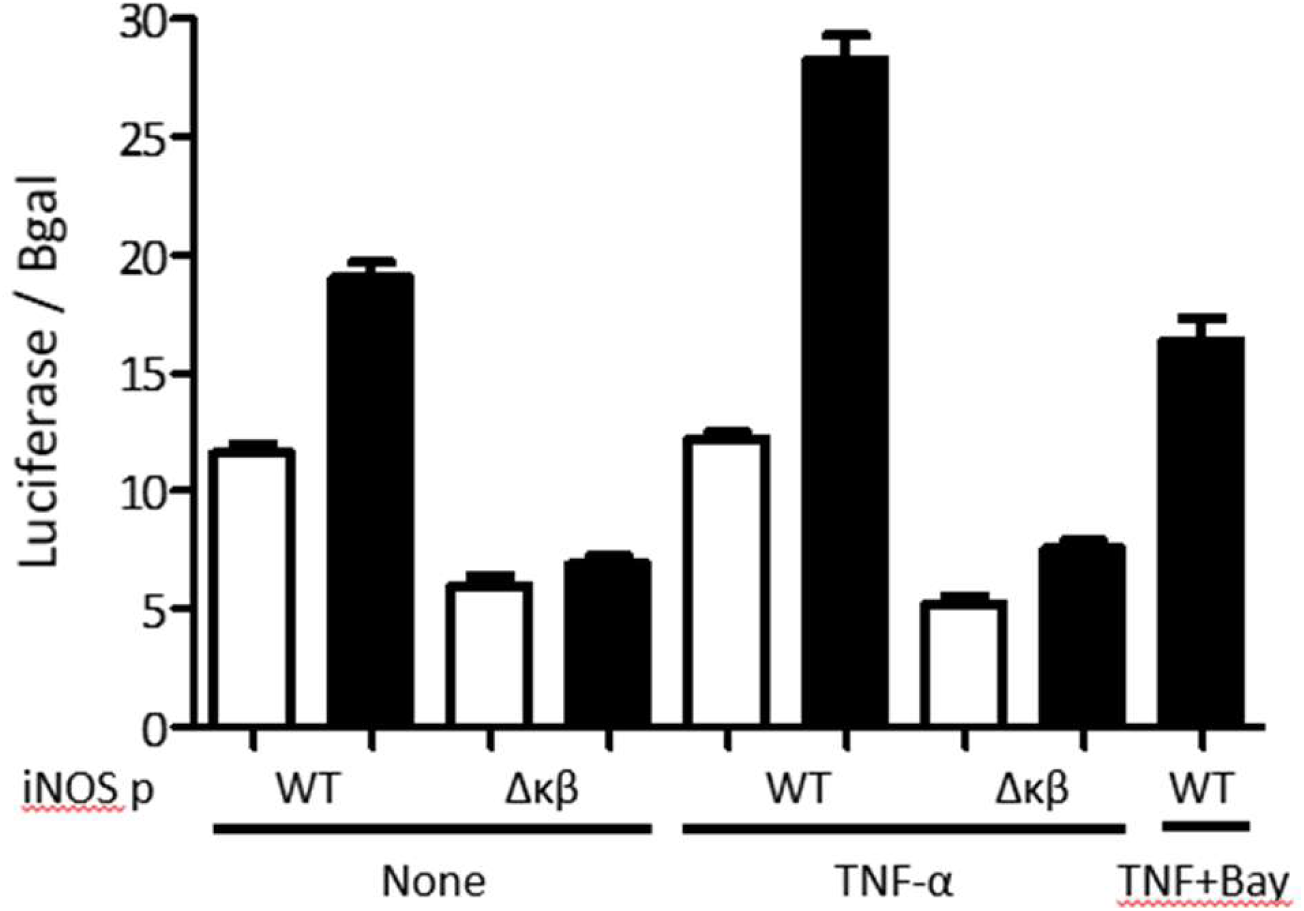
Effect of TNF-α stimulation on iNOS promoter transcriptional activity in cells expressing or no PARP-1. (**a**) PARP-1 levels in PARP-1 KO and BC HEK 293T cells. α-tubulin was detected as a loading control. (**b**) β-galactosidase activity-normalized luciferase levels in PARP-1 KO and BC HEK 293T cells transfected with a β-galactosidase expression plasmid and the iNOSpWT-luciferase or the iNOSpΔNFκB-luciferase reporter plasmid, and then treated with TNF-α or TNF-α and BAY. Data in (**b**) correspond to one experiment done in triplicate that is representative of two independent experiments performed in triplicate.

In correspondence with previous reports (47, 48), the basal expression of the iNOSpWT-luciferase plasmid was 1.6 folds lower in HEK 293T PARP-1 KO than in BC cells (Fig. 4b). In contrast, the basal expression of iNOSpΔNFκB-luciferase was similar in both cell lines but 1.9-fold lower in KO cells and 2.7-fold lower in BC cells compared to the expression driven by iNOSpWT (Fig. 4b). Furthermore, TNF-α activated iNOSpWT-luciferase in the HEK 293T PARP-1 BC cells (1.5 folds) but failed to stimulate this promoter in the PARP-1 KO cells. Furthermore, TNF-α treatment also failed to activate iNOSpΔNFκB-luciferase in both PARP-1 KO and BC HEK 293T cells (Fig. 4b). The activating effect of TNF-α on iNOSpWT-luciferase was entirely inhibited by BAY in PARP-1 BC cells (Fig. 4b), indicating the NF-κB-dependency of this observation.

### PARP-1 does not affect HIV-1 LTR silencing

PARP-1 has been proposed as a required cofactor for HIV-1 integration in the centromeric region (50). HIV-1 integration in centromeric region is disfavored (51, 52) and has been associated to the latent reservoir (53, 54). Therefore, we evaluated the temporal stability of gene expression of the HIV-1 provirus in SUPT1 cells expressing or no PARP-1. Hluc-infected SUPT-1 PARP-1 KO and BC cells were cultured for one month and luciferase expression was measured. Notoriously, HIV-1 gene expression was similar in cells expressing or no PARP-1 after prolonged cell culture (Fig. 5a), suggesting an equivalent tendency of HIV-1 proviruses to gene silencing in these cells.

**Figure 5.**
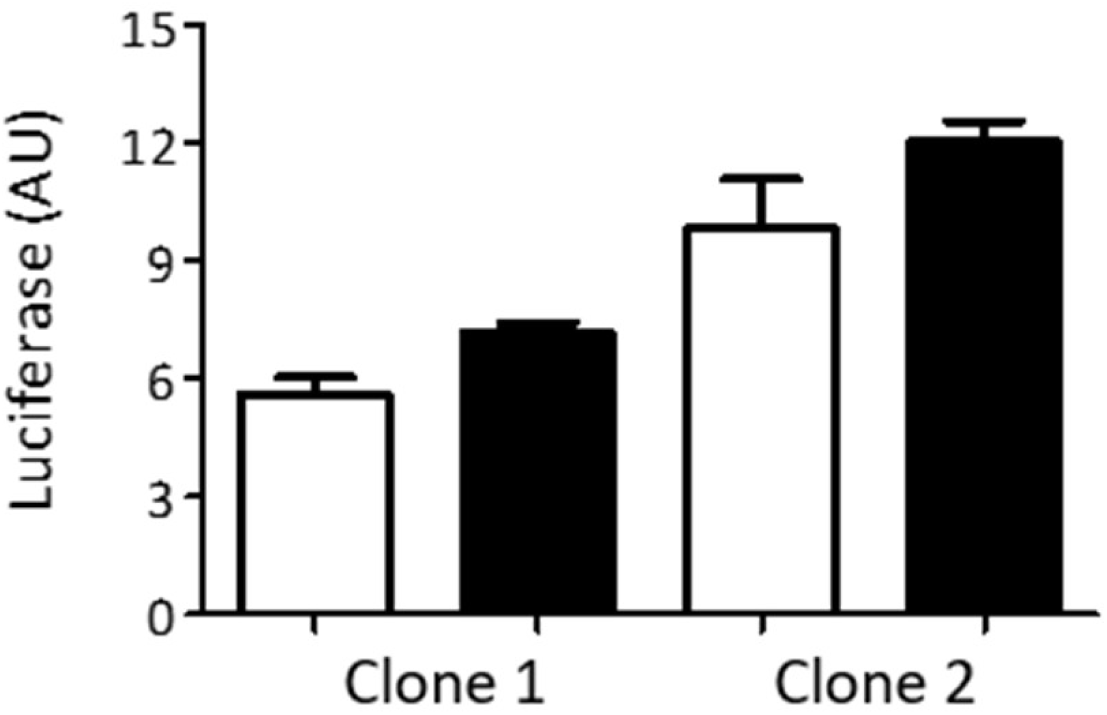
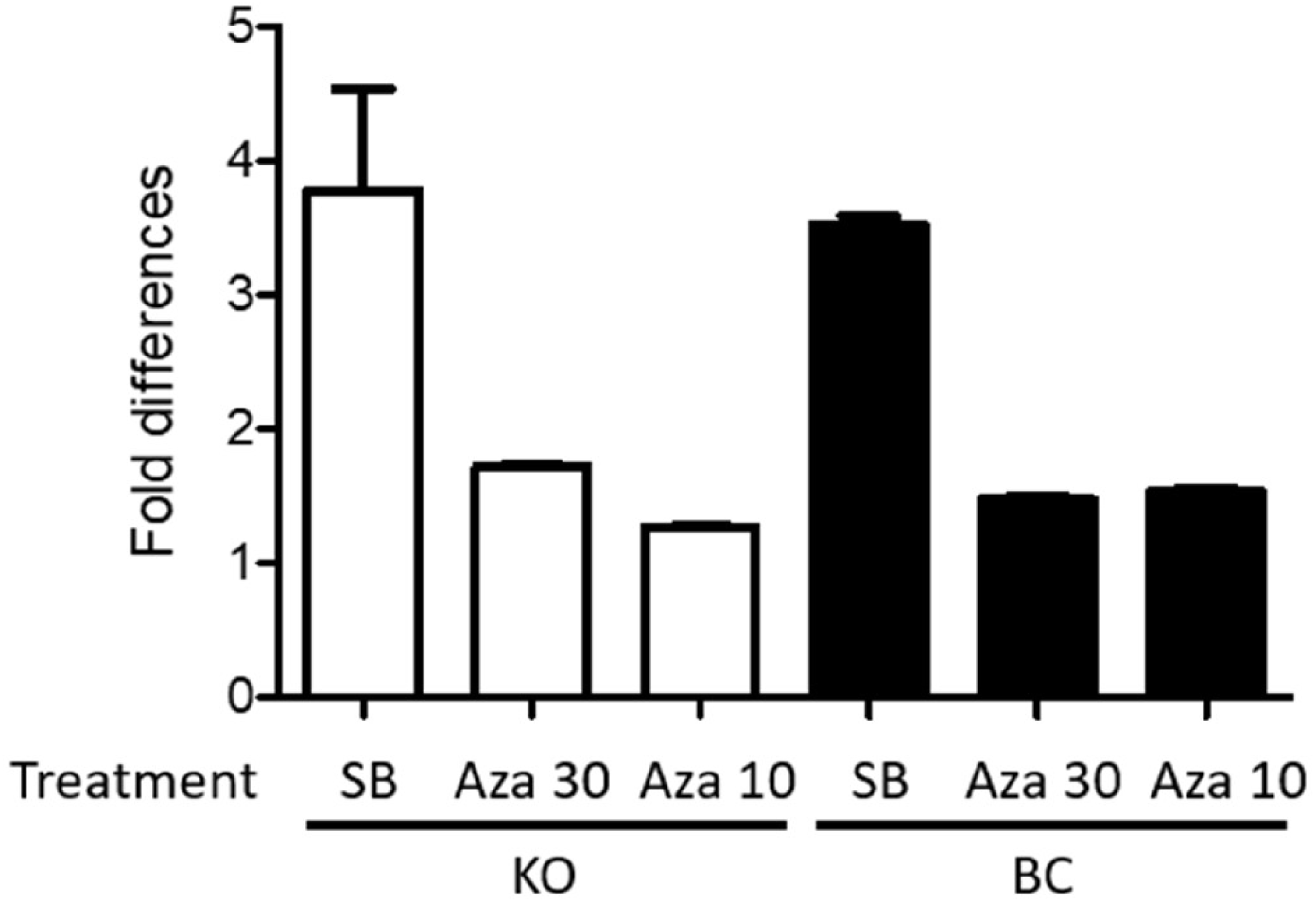
Effect of PARP-1 on HIV-1 provirus transcriptional silencing. (**a**) Luciferase levels in Hluc-infected PARP-1 KO and BC SUP-T1 cells cultured for one month. (**b**) Fold differences of ATP-normalized luciferase levels in cells characterized in panel (**a**) upon 24 or 36 hrs stimulation with sodium butyrate (5 uM) or 5-Azacytidine (30 and 10 uM), respectively. Data in Figure 5 correspond to one experiment conducted in triplicate, and is representative of two independent experiments.

Furthermore, these long-term infected cells were stimulated with sodium butyrate (SB, 5 mM) or 5-Azacytidine (Aza, 30 and 10 uM), known epigenetic modulators that activate silenced HIV-1 proviruses (55). ATP was measured in the treated cells to verify the preservation of the cell viability upon treatment, and ATP values were used to normalize the luciferase activity in these cells. SB was more potent activator than Aza in both cell lines, and both compounds similarly enhanced HIV-1 gene expression in PARP-1 KO and BC SUPT1 cells (Fig. 5b), indicating that these compounds activate HIV-1 gene expression independently of PARP-1. Furthermore, this data suggested that the size of the HIV-1 silenced reservoir susceptible to reactivation with these epigenetic modifiers is similar in both cell lines.

## DISCUSSION

NF-κB signaling is critical in the expression of inflammatory and immune genes (17–22), and in the transcriptional activity of the HIV-1 promoter during latency reactivation and active viral replication. Therefore, HIV-1 latency reactivation agents acting through NF-κB signaling are expected to trigger important inflammatory and immune reactions, limiting their clinical value. PARP-1 is required for NF-κB-mediated induction of inflammatory and immune genes (17–22). However, the role of PARP-1 in HIV-1 gene expression is debatable, and its requirement for NFκB-induced HIV-1 gene expression is ill-defined.

Our results indicate that PARP-1 is dispensable for basal or NF-κB-induced HIV-1 gene expression in a human CD4+ T cell line. These findings contradict reports indicating a positive or negative role of PARP-1 in HIV-1 transcription. In non-infected cells, PARP-1 has been reported to enhance HIV-1 LTR-mediated reporter gene expression under basal conditions (33) and upon stimulation with NF-κB-inducers (18, 30, 31, 34). In contrast with this positive role in HIV-1 transcription, PARP-1 has also been reported to negatively regulate the HIV-1 promoter when evaluated using LTR-reporter systems (56–59). These contradictions highlight important differences in the regulation of the HIV-1 LTR promoter when presented as a plasmid or as a provirus.

Our data in CD4+ T cells are also in contradiction with findings in cells of myeloid origin. PARP-1 inhibition was found to decrease HIV mRNA levels in the chronically infected human pro-monocyte U1 cell line stimulated with the NF-κB-inducer, PMA (30). This contradiction adds further evidence to the previously described differences in HIV-1 promoter activity in T cells and macrophages (60, 61).

In conclusion, we found that PARP-1 does not influence NF-κB-dependent HIV-1 gene expression in a human CD4+ T cell line. This is in marked contrast to the required role of PARP-1 in TNF-α-induced, NFκB-dependent activation of pro-inflammatory gene promoters (19, 27, 42, 43). The differential role of PARP-1 on HIV-1 and pro-inflammatory gene promoters illustrates the reported exquisite specificity of NF-κB signaling (62, 63). That is, NF-κB allows, for the same stimulus, to differentially modulate promoters with different requirements of transcription factors, co-activators, and co-repressors. Therefore, due to differences in promoter architecture, PARP-1 could play unique roles in NF-κB-dependent activation of HIV-1 LTR and pro-inflammatory gene promoters. Our findings further suggest that PARP-1 antagonism could reduce the pro-inflammatory effects of HIV-1 latency reactivation agents acting via NF-κB.

## MATERIALS AND METHODS

### PARP-1 knockout (KO) and backcomplemented (BC) cell lines

The generation and characterization of PARP-1 KO and backcomplemented cell lines was described in (39). Briefly, SUP-T1 and HEK 293T cells were transduced with an HIV-1-derived viral vector expressing a zinc-finger nuclease targeting PARP-1, and selected in the presence of puromycin. The lack of PARP-1 expression was verified by immunoblot, and single-cell KO clones were selected. To generate the backcomplemented counterpart, these clones were transduced with a Murine Leukemia Virus-derived viral vector expressing PARP-1, and then selected in the presence of G418. PARP-1 re-expression was verified by immunoblot.

### Generation of lentiviruses

The replication-defective, HIV-1 reporter viruses were produced by calcium-phosphate transfection of HEK 293T as previously described (64). Briefly, cells were co-transfected with the HIV-1 transfer plasmid (15 ug), the HIV-1 packaging plasmid pCMVΔR8.91 (15 ug), and the plasmid pMD.G (5 ug) encoding the Vesicular Stomatitis Virus glycoprotein G. The HIV-1 reporter NLENG1-ES-IRES (65) is *env* deleted and expresses LTR-driven eGFP from the *nef* slot, and Nef from an IRES. The HIV-1 reporter Hluc expresses LTR-driven luciferase from the *nef* slot and contains a large deletion in *env* (40). This virus also lacks the expression of Vpr and Nef. Hluc_Δκβ_ was derived from Hluc by mutating the two LTR NF-κB binding sites. To this end, PCR-mediated mutagenesis was performed with reverse primer EF3 (5’- ggaaagtagattgtagcaagctcgatgtcagcagttc-3’, target sequence 5’- gaactgctgacatcgagcttgctacaa**<u>tct</u>**actttcc-3’) and forward primer EF4 (5’- gctg**<u>tct</u>**actttccagggaggcgtggcctgggcgggactggggag-3’) using Phusion Site-Directed Mutagenesis Kit (Thermo Scientific catalog # F-541), as described in (64). The mutations introduced have been reported to ablate the two copies of the enhancer κB elements implicated in NF-κB binding to the LTR (46), and are indicated in the primers with bolded and underlined font.

### Single-round infection analysis

PARP-1 KO and BC SUP-T1- and HEK 293T-derived cell lines were plated at a density of 01.X10^6^ cells in 500 µl of culture medium per well in 24-well plates and infected with the non-replicating viruses. Cells infected with viruses expressing luciferase were typically harvested four days after infection and analyzed for luciferase activity with a luminescence kit (Bright-Glo™ Luciferase Assay System, Promega, E2620) as described in (64). In some experiments, ATP was measured in the same samples using the CellTiter-Glo® Luminescent Cell Viability Assay (Promega, G7570) to normalize for cell number and viability. Cells infected with the HIV-1 reporter expressing eGFP were analyzed by flow cytometry four days after infection.

### Effect of different stimuli on HIV-1 infection

HIV-1-infected SUP-T1 PARP-1 KO and BC cells (2×10^5^ cells / 300 µl) were subjected to different stimuli for three days and then luciferase levels were measured. The stimuli were TNF-α (10 ng/ml), anti-CD3/CD28 immunobeads (1 bead per cell, Dynabeads® Human T-Activator CD3/CD28 for T Cell Expansion and Activation, catalog number 11161D), and PMA (10 ng/ml). In some experiments, cells were treated with the NF-κB signaling inhibitor BAY-11-7082 (3 µM) one hour before TNF-α stimulation. Then the inhibitor was kept for the entire duration of the experiment.

Long-term HIV-1-infected SUP-T1 PARP-1 KO and BC cells (2×10^5^ cells / 300 µl) were treated with sodium butyrate (5 mM; B5887; Sigma) for 24 h or with 5-azacytidine (30 and 10 µM; A2385; Sigma) for 36 h. Following these treatments, the cells were analyzed for luciferase and ATP levels. Luciferase levels were normalized to ATP levels to account for any potential effect of these treatments on cell viability.

### Effect of TNF-α on the activity of the iNOS promoter

PARP-1 KO and BC HEK 293T cells were calcium-phosphate co-transfected with a CMV-driven β-galactosidase expression plasmid [pCMV-β-gal, (66)] and iNOSpWT-luciferase or iNOSpΔNFκB-luciferase reporter plasmid (49). Transfected cells were subjected to different treatments, and β-galactosidase and luciferase were measured three days later. β-galactosidase was measured with the Beta-Glo assay system (Promega, E4720). Luciferase levels were normalized to β-galactosidase activity to control for transfection efficiency.

The iNOS reporters were previously described (49). The reporter iNOSpWT-luciferase contains a 1516-base pair fragment that includes 1485 and 31 nucleotides upstream and downstream, respectively, of the transcription start site of the mouse iNOS gene. The NF-κB site (−85 to −83 nucleotides) was mutated (GGG to CTC) in iNOSpWT-luciferase to generate iNOSpΔNFκB-luciferase.

## Acknowledgments

This work was supported by Grant Number SC1GM115240 to ML from the National Institute of Allergy and Infectious Diseases (NIAID), National Institutes of Health (NIH). We thank the Biomolecule, Genomic Analysis and the Cellular Characterization, and Biorepository Core Facilities for technical help. These core facilities are supported by a Research Centers in Minority Institutions program grants 5G12MD00759 and 2U54MD007592 to the Border Biomedical Research Center in UTEP from the National Institute on Minority Health and Health Disparities, a component of NIH.

We thank David N. Levy (New York University, Dental Center) for providing the HIV-1 reporter plasmid NLENG1-ES-IRES, Mark A. Perrella (Harvard Medical School) for providing the iNOSpΔNFκB- and iNOSpWT-luciferase reporter plasmids, and Elias Farran (UTEP) for generating the Hluc_Δκβ_ mutant.

